# Grey Matter Age Prediction as a Biomarker for Risk of Dementia: A Population-based Study

**DOI:** 10.1101/518506

**Authors:** Johnny Wang, Maria J. Knol, Aleksei Tiulpin, Florian Dubost, Marleen de Bruijne, Meike W. Vernooij, Hieab H.H. Adams, M. Arfan Ikram, Wiro J. Niessen, Gennady V. Roshchupkin

**Affiliations:** Department of Medical Informatics, Erasmus MC University Medical Center, Rotterdam, The Netherlands; Mechanical Engineering, Faculty of Mechanical, Maritime and Materials Engineering, Delft University of Technology, Delft, The Netherlands; Department of Epidemiology, Erasmus MC University Medical Center, Rotterdam, The Netherlands; Research Unit of Medical Imaging, Physics and Technology, University of Oulu, Oulu, Finland; Department of Computer Science, University of Copenhagen, Copenhagen, Denmark; Department of Radiology and Nuclear Medicine, Erasmus MC University Medical Center, Rotterdam, The Netherlands; Imaging Physics, Faculty of Applied Sciences, Delft University of Technology, Delft, The Netherlands

**Keywords:** Deep Learning, age prediction, biomarker, dementia, magnetic resonance imaging, brain, voxel-based morphometry, survival analysis

## Abstract

**Key Points:** *Question:* Is the gap between brain age predicted from MRI and chronological age associated with incident dementia in a general population of Dutch adults?

*Findings:* Brain age was predicted using a deep learning model, using MRI-derived grey matter density maps. In a population based study including 5496 participants, the observed gap was significantly associated with the risk of dementia.

*Meaning:* The gap between MRI-brain predicted and chronological age is potentially a biomarker for dementia risk screening.

**Abstract:** *Importance:* The gap between predicted brain age using magnetic resonance imaging (MRI) and chronological age may serve as biomarker for early-stage neurodegeneration and potentially as a risk indicator for dementia. However, owing to the lack of large longitudinal studies, it has been challenging to validate this link.

*Objective:* We aimed to investigate the utility of such a gap as a risk biomarker for incident dementia in a general Dutch population, using a deep learning approach for predicting brain age based on MRI-derived grey matter maps.

*Design:* Data was collected from participants of the cohort-based Rotterdam Study who underwent brain magnetic resonance imaging between 2006 and 2015. This study was performed in a longitudinal setting and all participant were followed up for incident dementia until 2016.

*Setting:* The Rotterdam Study is a prospective population-based study, initiated in 1990 in the suburb Ommoord of in Rotterdam, the Netherlands.

*Participants:* At baseline, 5496 dementia- and stroke-free participants (mean age 64.67±9.82, 54.73% women) were scanned and screened for incident dementia. During 6.66±2.46 years of follow-up, 159 people developed dementia.

*Main outcomes and measures:* We built a convolutional neural network (CNN) model to predict brain age based on its MRI. Model prediction performance was measured in mean absolute error (MAE). Reproducibility of prediction was tested using the intraclass correlation coefficient (ICC) computed on a subset of 80 subjects. Logistic regressions and Cox proportional hazards were used to assess the association of the age gap with incident dementia, adjusted for years of education, ApoEε4 allele carriership, grey matter volume and intracranial volume. Additionally, we computed the attention maps of CNN, which shows which brain regions are important for age prediction.

*Results:* MAE of brain age prediction was 4.45±3.59 years and ICC was 0.97 (95% confidence interval CI=0.96-0.98). Logistic regression and Cox proportional hazards models showed that the age gap was significantly related to incident dementia (odds ratio OR=1.11 and 95% confidence intervals CI=1.05-1.16; hazard ratio HR=1.11 and 95% CI=1.06-1.15, respectively). Attention maps indicated that grey matter density around the amygdalae and hippocampi primarily drive the age estimation.

*Conclusion and relevance:* We show that the gap between predicted and chronological brain age is a biomarker associated with risk of dementia development. This suggests that it can be used as a biomarker, complimentary to those that are known, for dementia risk screening.

## 1. Introduction

The human brain continuously changes throughout the entire lifespan. These changes partially reflect a normal aging process and are not necessarily pathological^1^. However, neurodegenerative diseases, including dementia, also affect brain structure and function^2,3^. Therefore, a better understanding and modeling of normal brain aging can help to disentangle these two processes and improve the detection of early-stage neurodegeneration.

Age prediction models based on brain magnetic resonance imaging (MRI) are a popular trend in neuroscience^4–7^. The difference between predicted and chronological age is thought to serve as an important biomarker reflecting pathological processes in the brain. Several recent studies showed the relation between accelerated brain aging and various disorders, such as Alzheimer’s disease (AD), schizophrenia, epilepsy or diabetes^7–9^.

In recent years, convolutional neural networks (CNN) have become the methodology of choice for analyzing medical images. These models are able to learn complex relations between input data and desired outcomes. Recent studies were able to demonstrate that CNN models can outperform complex machine learning models in brain MRI-based age prediction^5,6^.

Although cross-sectional studies have suggested that the gap between predicted and chronological age may serve as a biomarker for dementia diagnosis, it remains unclear whether this is also the case for the years preceding dementia diagnosis^5,7^. Longitudinal studies examining the link between such a gap and incident dementia are lacking and are crucial for validation of this biomarker for early-stage neurodegeneration detection. Using a deep learning (DL) model, we investigated the association of the grey matter (GM) age gap with incident dementia in a large population-based sample of middle-aged and elderly subjects.

## 2. Methods

### 2.1 Study Population

Data was acquired from the Rotterdam Study, an ongoing population-based cohort study among the inhabitants of Ommoord, a suburb of Rotterdam, the Netherlands^10^. The cohort started in January 1990 (n=7983) and was extended in February 2000 (n=3011) and February 2006 (n=3932). Follow-up examinations take place every 3 to 4 years. MRI was implemented in 2005, and 5912 persons scanned until 2015 were eligible for this study. We excluded individuals with incomplete acquisitions, scans with artifacts hampering automated processing, participants with MRI-defined cortical infarcts and participants with dementia or stroke at the time of scanning (**Supplementary figure 1**). This resulted in 5656 subjects to be included in this study. The Rotterdam Study has been approved by the Medical Ethics Committee of the Erasmus MC and by the Ministry of Health, Welfare and Sport of the Netherlands, implementing the Wet Bevolkingsonderzoek ERGO (Population Studies Act: Rotterdam Study). All participants provided written informed consent to participate in the study and to obtain information from their treating physicians.

### 2.2 Image processing

A 1.5 tesla GE Signa Excite MRI scanner was used to acquire multi-parametric MRI brain data, as previously reported^10^. Voxel-based morphometry (VBM) was performed according to an optimized VBM protocol as was previously described^11,12^. First, all T1-weighted images were segmented into supratentorial GM, white matter (WM), and cerebrospinal fluid (CSF) using a previously described *k*-nearest neighbor algorithm, which was trained on six manually labeled atlases^13^. FMRIB’s Software Library (FSL) software was used for VBM data processing^14^. All GM density maps were non-linearly registered to the standard Montreal Neurological Institute (MNI) GM probability template, with a 1×1×1 mm^3^ voxel resolution.

A spatial modulation procedure was used to avoid differences in absolute GM volume due to the registration. This involved multiplying voxel density values by the Jacobian determinants estimated during spatial normalization. We did not apply smoothing. While VBM smoothing procedures increase the signal to noise ratio, they can affect the features which the network learns from GM.

Intracranial volume (ICV) estimates were obtained by summing total GM, WM and CSF volumes.

#### Dementia assessment

All participants were monitored for dementia at baseline and following visits to the study center using the Mini-Mental State Examination (MMSE) and the Geriatric Mental State (GMS) organic level. Further investigation was initiated for participants who scored lower than 26 for their MMSE or above 0 for their GMS^15^. Additionally, the entire cohort was continuously checked for dementia through electronic linkage between the study center and medical records from general practitioners and the regional institute for outpatient mental health care. Available information on cognitive testing and clinical neuroimaging was used when required for diagnosis of dementia subtype. Final diagnosis was established by a consensus panel led by a consultant neurologist, according to a standard criteria for dementia (using the Third Revised Edition of the Diagnostic and Statistical Manual of Mental Disorders (DSM-III-R))^16,17^. Until January 1^st^ 2016, 92% of the potential person-time follow-up was complete. Participants were censored at date of dementia diagnosis, death or loss to follow-up, or at January 1st 2016, whichever came first. Of 5496 subjects included in this analysis, 159 developed dementia within 10 years of follow-up (mean follow-up time 4.34±2.25 years).

#### Other measurements

ApoEε4 carriership was determined using a polymerase chain reaction (PCR) on coded deoxyribonucleic acid (DNA) samples. If these values were missing, Haplotype Reference Consortium (HRC) imputed genotype values for rs7412 and rs429358 were used to define the ApoEε4 carrier status. Measurements on more characteristics are described in **Supplementary Methods 1**.

### 2.3 Deep Learning model

A full description of the applied DL model is presented in the **Supplementary Methods 2**. Briefly, a DL model takes a set of inputs and respective outputs from a training set and finds an optimal non-linear relation between them. A CNN is a class of DL techniques which takes in multi-dimensional images as model input. These networks are generally used with a variety of different techniques and algorithms, which together define how the model optimizes the input-output relationship^18,19^. We describe this in details in the model architecture.

Our 3-dimensional (3D) regression CNN model is designed to predict brain age using 3D GM density maps from VBM as input. It is inspired by ConvNet^20^ and Deep CNN^19^, as shown in **Supplementary Figure 2**. Besides GM brain images, we provide information about the sex of the subject. This allows the network to adjust for GM differences between male and female subjects.

The dataset, excluding subjects with incident dementia, was randomly split into three sets: training (3688 subjects), validation (1099 subjects) and test (550 subjects). Subjects with incident dementia (159 subjects) were put in a fourth independent dataset. The CNN was trained using the training set as described in **Supplementary Methods 3**. For training we used all available scans for each subjects. Prediction accuracy was assessed on the test set. Model accuracy was measured based on the absolute gap, or mean absolute error (MAE) of prediction, i.e. the difference between model output and real chronological age (gap = age_brain, predicted_ – age_chronological_).

#### Attention mapping

We retrieved attention maps from the trained networks using Gradient-weighted Class Activation Mapping (Grad-CAM)^21^. Attention maps show which areas on subject GM image are more important for age prediction. Attention map intensity values were normalized to range 0-1, where 1 indicates the value for areas most associated with the network’s decision. We expanded the Grad-CAM visualization technique to a 3-dimensional space.

Attention maps were computed for every individual. Since all brain images were registered to the same template space, a global average voxel-wise attention map could be made over attention maps of all subjects to obtain a global attention map for the age prediction network.

We computed the change in attention map over age per voxel, to investigate the change in regions predictive for brain age between age groups. To this end, for each voxel, a linear regression from age to attention map value was performed, resulting in a line of which the slope represents the increase in attention map value with age for the given voxel.

### 2.4 Statistical analysis

Reproducibility of the CNN age prediction was quantified using the intraclass correlation coefficient (ICC(3,1)), computed on a subset of 80 persons out of the test set who were scanned twice with a time interval of one to nine weeks^22^.

In order to be able to compare our findings with previous studies, logistic regression models and Cox proportional hazards models were used to assess the association between the age gap and the incidence of dementia. We adjusted the regression models for biomarkers, which are known for their relation with dementia: age and sex (model I); additionally GM volume and ICV (model II); and years of education and APOEε4 carriership (model III)^23,24^. The logistic regression model used the occurrence of dementia-development during follow-up as output. The proportional hazards and linearity assumption were met for the Cox proportional hazards models. Python and R were used to perform the statistical analyses^25–28^.

## 3. Results

The study population characteristics are described in **Table 1**. The algorithm was trained and validated on random subsets of subjects with mean age 66.09±10.76 years and 55% females; and mean age 64.84±9.69 years and 54% females, respectively. The following results are reported for the test set (mean age 64.85±10.82 years and 55% females).

**Table 1.** Characteristics of data sets derived from the population-based Rotterdam Study.

|  | Train | Validation | Test** | Incident dementia** |
| --- | --- | --- | --- | --- |
| N<sub>subj</sub> | 3688 | 1099 | 550 | 159 |
| N<sub>img</sub> | 5865 | 2353 | 550 | 159 |
| Mean age* (years±sd) | 66.09±10.76 | 64.84±9.69 | 64.85±10.82 | 77.33±7.15 |
| Sex proportion* (female/male) | 0.55/0.45 | 0.54/0.46 | 0.55/0.45 | 0.58/0.42 |
| Education* (years±sd) | 12.64±3.89 | 12.63±3.81 | 12.58±4.00 | 11.43±3.57 |
| GM volume* (liters±sd) | 0.60±0.06 | 0.60±0.06 | 0.60±0.06 | 0.55±0.05 |
| ICV* (liters±sd) | 1.48±0.16 | 1.47±0.16 | 1.48±0.16 | 1.45±0.17 |
| ApoE4 carriership* (0/1/2) | 0.72/0.26/0.02 | 0.72/0.25/0.02 | 0.74/0.23/0.03 | 0.57/0.36/0.06 |
| Follow-up time* (years±sd) | 5.42 ±2.81 | 4.93±2.80 | 6.68±2.29 | 4.29±2.26 |

\* Values are based on N<sub>img</sub>.
\*\* Selection only includes baseline image of subjects.
*Abbreviations:* number of subjects (N<sub>subj</sub>); number of images (N<sub>img</sub>); grey matter (GM); intracranial volume (ICV)

### 3.1 Network performance

The overall performance measured on the test set was MAE=4.45±3.59 years (**Figure 1**), with a correlation between chronological and predicted brain age of 0.85 (p-value=4.76×10^−156^). A reproducibility score of ICC=0.97 (95% confidence interval CI 0.96-0.98) was achieved. No significant difference in prediction was found between male and female subjects (p-value=0.34), detailed numbers are provided in **Supplementary Text 1**.

**Figure 1.**
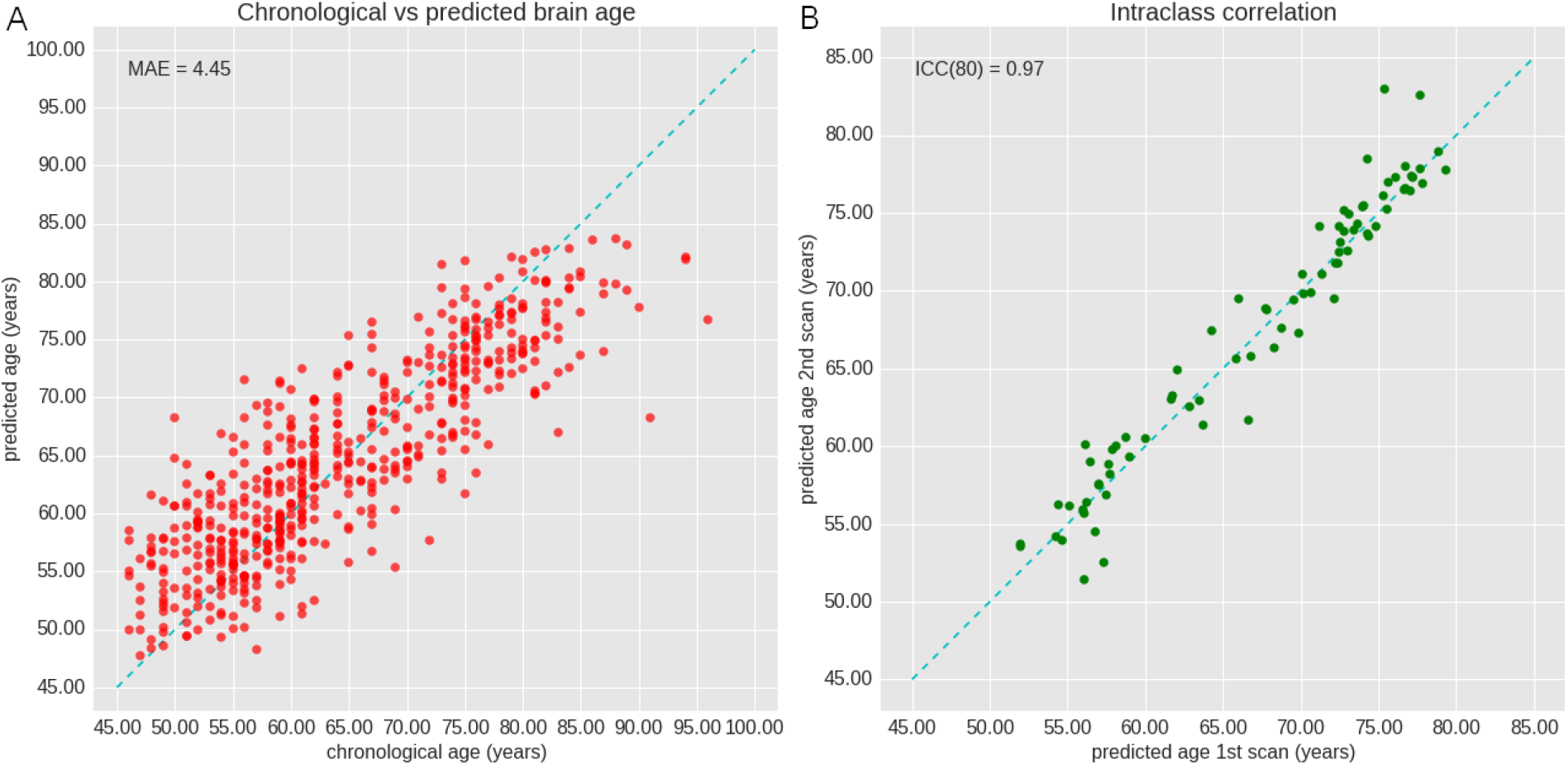
Performance of CNN on test dataset. **(A)** The plot depicts chronological age (x-axis) and brain-predicted age (y-axis) with mean absolute error (MAE). The dashed line indicates the ideal case x=y. **(B)** The figure shows reproducibility of the CNN performance. Scan 1 and 2 are taken with one to nine weeks interval. The dashed line indicates a perfect reproducibility and consistent predicting of the network.

#### Attention map

**Supplementary Figure 5** shows the global attention map of the test set, indicating the areas contributing to age prediction in bright color, as well as the increase of attention map values over age. We found that the amygdala and hippocampus are not only important for predicting brain age, but that these regions also grow more important with increasing chronological age, which is shown in **Supplementary Figure 5B**. A quantitative analysis per brain region is presented in **Table 2**, which shows that highest mean intensities were computed for the nucleus accumbens (0.89) and amygdala (0.71). Highest intensity quintiles were computed for the nucleus accumbens (0.99), amygdala (0.98) and subcallosal area (0.98).

**Table 2.**
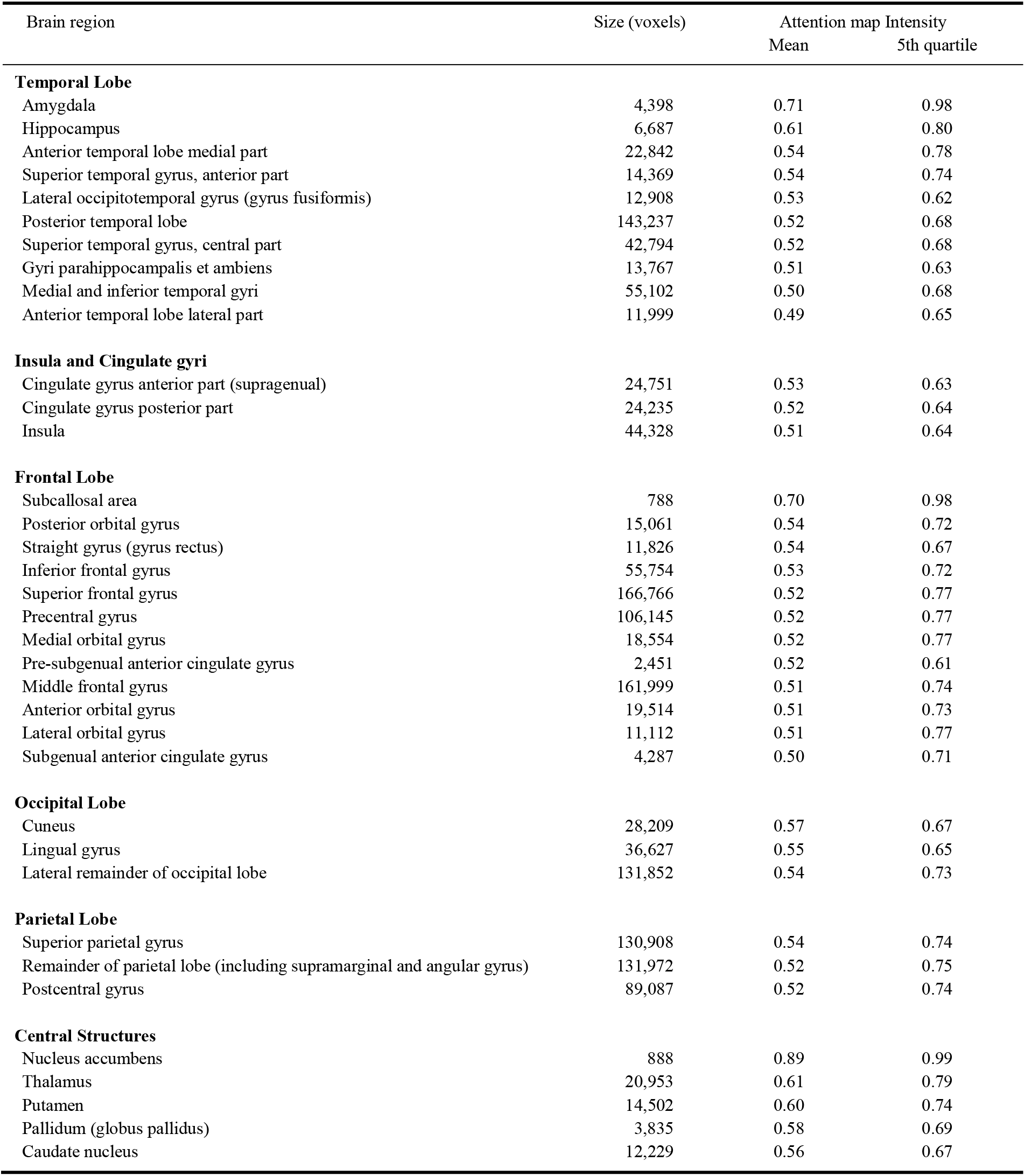
Quantitative analysis of the attention map per brain region. Mean and fifth quintile (lower boundary) of attention map intensity per brain region are listed. Brain regions are grouped by lobes.

### 3.2 Logistic regression

We computed a logistic regression for the three models, as shown in **Table 3**. The age gap was significantly associated with dementia incidence while age, sex, education years, GM and ICV volume and the ApoEε4 allele carriership were included in the model, with model III: odds ratio OR=1.11 (95% CI 1.05-1.16) per year age gap.

**Table 3.** Association of gap between brain age and chronological age with incident dementia assessed by logistic regression and Cox proportional hazards models, both in the total study sample and in a subsample with a minimum follow-up time of 5 years.

| Model | Logistic Regression |  |  | Cox Regression |  |  |
| --- | --- | --- | --- | --- | --- | --- |
|  | n/N | OR (95% CI) | p-value | n/N | HR (95% CI) | p-value |
| Total sample |  |  |  |  |  |  |
| Model I | 159/1808 | 1.15 (1.10-1.20) | $2.67 \times 10^{-10}$ | 159/1808 | 1.15 (1.11-1.20) | $1.02 \times 10^{-12}$ |
| Model II | 154/1790 | 1.11 (1.06-1.16) | $2.57 \times 10^{-5}$ | 154/1790 | 1.11 (1.07-1.16) | $4.59 \times 10^{-7}$ |
| Model III | 150/1714 | 1.11 (1.05-1.16) | $4.80 \times 10^{-5}$ | 150/1714 | 1.11 (1.06-1.15) | $1.23 \times 10^{-6}$ |
| Sample follow-up time > 5 years |  |  |  |  |  |  |
| Model I | 62/1366 | 1.11 (1.04-1.18) | $1.26 \times 10^{-3}$ | 62/1366 | 1.13 (1.06-1.20) | $1.38 \times 10^{-4}$ |
| Model II | 60/1352 | 1.09 (1.02-1.16) | $1.43 \times 10^{-2}$ | 60/1352 | 1.10 (1.03-1.17) | $3.20 \times 10^{-3}$ |
| Model III | 58/1305 | 1.09 (1.01-1.16) | $2.08 \times 10^{-2}$ | 58/1305 | 1.09 (1.02-1.17) | $7.24 \times 10^{-3}$ |

Model I: age + sex.
Model II: model I + grey matter volume + intracranial volume.
Model III: model II + years of education + APOEε4 carrier status.
Abbreviations: confidence interval (CI); odds ratio (OR); hazard ratio (HR); number of cases (n); total number of participants (N).

### 3.3 Survival analysis

As shown in **Table 3** and **Figure 2**, the age gap was significantly associated with the incidence of dementia, with model III hazard ratio HR=1.11 (95% CI 1.06-1.15) per year age gap. These associations were similar in a subsample with a follow-up time for indecent dementia of more than 5 years, model III HR=1.09 (95% CI 1.02-1.17) per year age gap.

**Figure 2.**
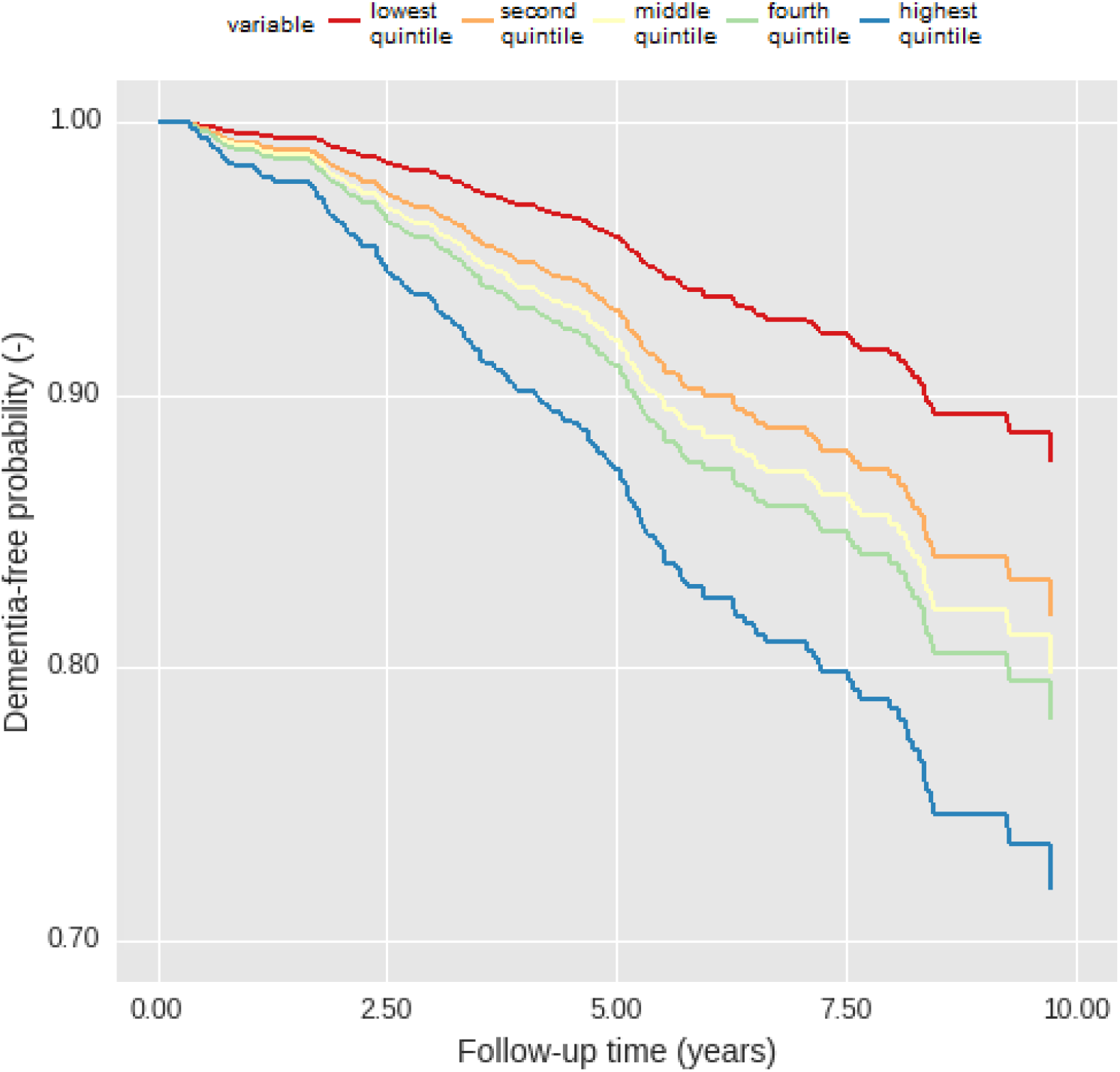
Adjusted survival curves for dementia-free probability by age gap. Dementia-free probability is presented over time for participants with different age gap values, divided into quintiles. Lower gap values correspond to chronological ages surpassing brain age, whereas higher gap values correspond to chronological ages that are lower than the brain age. Plots are based on Cox proportional hazards models, adjusted for age, sex, total grey matter volume, intracranial volume, years of education and ApoEε4 carriership status, using a marginal approach.

### 3.4 Gap-associated features

**Supplementary Table 1** shows a list of features that can affect the brain pathology and may be associated with the gap^9^. Significantly lower values were found for GM volume in the highest quintile. However, systolic blood pressure and mild cognitive impairment were already only nominally significant, after Bonferroni correction.

## 4 Discussion

In a large sample of community-dwelling middle-aged and older adults, using a DL model for brain age prediction on MRI-derived grey matter tissue density, we found that the gap between predicted brain age and chronological age was related to an increased risk of dementia, independent of standard established risk factors for dementia.

Our trained CNN model showed a similar performance in age prediction compared to previous studies that use a multimodal data model^5^ and DL-based approach^6^, which achieved performances of MAE=4.29 and MAE=4.16, respectively. Previous studies looked cross-sectionally^5,6^ at the association of the age gap and dementia occurrence, while in the current study we evaluated associations in longitudinal data. As non-reversible pathological changes already occur years prior to diagnosis, identifying early-stage biomarkers for dementia is of importance. The age gap has potential to be utilized alongside other clinical risk factors and biomarkers to separate the population into categories with sufficiently distinct degrees of risk to drive clinical or personal decision-making, e.g. dementia screening and informed life planning.

Moreover, we retrieved attention maps from the model, showing which brain regions are most important for age prediction, which also provides insights into processes in aging. Since specific regions were identified on which the model mainly focused, this suggests that the gap holds more specific information than global measures of GM volume when predicting brain age. This was further established by the association found between the gap and incident dementia, which remained significant after adjusting for total GM volume. Based on the attention maps the amygdala and hippocampus in particular proved to be more associated to age prediction, also increasing in attention map intensity with older subjects. This is in accordance to literature where significant negative associations between GM volume and age have been reported for these regions^2,23^. Atrophy of these two structures has also shown to be more prevalent in dementia patients, including years before diagnosis^29,30^. A more in depth evaluation of the attention map can be found in **Supplementary text 2**.

### Limitations

We were not able to perfectly predict the age for healthy subjects based only on MRI. We assume that due to biological similarity of the brain within a range of several years, there will always be an according level of uncertainty in the age prediction.

Furthermore, we excluded subjects with dementia and stroke while training the model, but there are a number of other factors which can influence overall or local GM volume and affect the age prediction and gap (**Supplementary Table 1**). Although only total GM volume differed significantly between subjects with a high versus a low gap, effect estimates of some features differed substantially. Further research is needed to investigate gap-associated features, which may explain gap differences. These features can also introduce bias, which may be solved by adding the information as a covariate to the model. This however requires the respective information on the subjects, which can make the method less accessible for general use.

Lastly, the current CNN model is incapable of handling unfamiliar datasets, limiting its practical use. A drawback of CNN’s is that the training data should be representative for the data for which the trained network is used. Thus limiting the generalizability of our method. However, this can be addressed by training models on more diverse or new datasets. It would therefore be interesting to extend this model to another dataset and validate its use in a different context

## 5. Conclusion

We showed that the gap between age predicted from brain MRI and chronological brain age is a biomarker associated with a risk of dementia development. DL visualization allows further investigation of the gap and neurodegeneration with respect to the human brain. This suggests that the age gap may be applicable for dementia risk screening, but there is still room for improvement in accuracy and for further research into the association between gap and dementia compared to other biomarkers.

## Acknowledgements

Mr. Aleksei Tiulpin was supported by KAUTE foundation.

## SUPPLEMENTARY MATERIAL

### Supplementary methods 1: Measurements of characteristics

Mild cognitive impairment (MCI) was assessed in individuals over the age of 60 years, for which both subjective and objective cognitive deficits were required. An objective cognitive deficit was based on a cut-off of 1.5 standard deviations below the Rotterdam Study age- and education-specific means in three cognitive domains, i.e. the memory, information processing speed and executive functioning domain. Subjective cognitive deficits were defined as having answered yes to any of six questions regarding difficulties in memory (difficulties finding words, or remembering plans) or daily functioning (difficulties managing finances, getting dressed, or using the phone).

Systolic and diastolic blood pressure was measured twice in the right arm in sitting position after five minutes of rest, of which the average was used. Body mass index (BMI) was defined as weight in kilograms (kg) divided by height in meters squared (m^2^). Participants were asked by interview whether they were a current or past smoker, which was used to define their smoking status. Glucose, total cholesterol and HDL cholesterol were measured in blood of the fasting state.

### Supplementary methods 2: Deep learning and convolutional neural networks

Deep learning techniques require a set of input and respective output to find and optimize a non-linear relation between the two. By providing data to a set of algorithms, the method is able to train a by the user designed model. Generally, the user designs the model architecture by selecting the model components. Subsequently, the machine learning method iteratively adjusts the model parameters according to that iteration’s trained model performance, to create an optimized model using backpropagation by supervised or unsupervised learning^1,2^. By letting the model itself choose which relevant features to extract from the input, deep learning facilitates the model to freely search the input-space and find the most important, possibly new, input features.

Convolutional neural networks (CNNs) are a subset amongst deep learning techniques. They allow multi-dimensional input images and inspect these inputs by scanning them for relevant information^3,4^. Deep learning and CNN models have been rising in popularity and have been actively studied in recent years, reaching state-of-the-art performances in many applications amongst which medical imaging^5–7^.

CNNs regard an image as a field of numerical values, view small portions of this image (receptive field) and perform multiplications with a weight-matrix (filter) to extract certain information (feature) from this portion. By inspecting the entire image using this filter in a grid-wise manner, the filter extracts specific information which is then saved to a new matrix or image (feature map). Repeating this process for the resulting feature maps, the network iteratively refines or searches for more information inside of the image that is relevant to the output.

These convolutional layers (CONV layers) are then typically combined with a variety of different techniques and algorithms that allow the network to appropriately extract the information from the input. Commonly used techniques are rectified linear units activation (ReLU), max-pooling layers (MP), fully connected layers (FC), batch normalization and dropout^4,7^.

### Supplementary methods 3: Network training

The CNN has been trained using the data from the training set of 3688 subjects. Here, over- and undersampling had been applied to the training set. Thus, effectively data of 3688 subjects was used out of 3848 available subjects available for the training set, to distribute the samples more evenly over the age range of the population (N_img,train_balanced_=8060 images, mean age 68.52±13.71sd). To avoid overfitting on the training set and to improve overall model performance, data augmentation was also applied during training^8^. Data augmentation included random small translations and mirroring in planes. We also used follow-up MRI scans of each subject as a ‘natural data augmentation’ technique.

The best model was selected based on its performance on the validation set. Here the performance is measured as the model accuracy based on the root mean squared error (RMSE) of the gap, as RMSE penalizes outliers more than MAE.

### Supplementary text 1: Sex covariate effect on CNN model performance

We can consider a split evaluation between male and female subjects. **Supplementary Figure 3** shows the network found no significant difference between the two groups (p=0.34). By including sex as a covariate, the covariate can reduce the difference in resulting age predictions between male and female subjects.

The trained model was able to reduce prediction error and correct for male and female biases observed in the image by the model. By including the additional input of sex, the model is able to prevent over- and under prediction for male and female ages, respectively, as shown in **Supplementary Figure 4**. Here we present the performance in gap on male and female subjects, both early adapted models were trained under the same training settings and used the exact same training and validation sample sets. The model that includes the additional input of the subject’s respective sex, was able to reduce the overall gap between male and female subjects to be insignificant (p-value=0.23). This also brought the mean gap for males and females closer to zero (one-sample t-test: p_male_=0.88 and p_female_=0.05).

### Supplementary text 2: Important regions attention map

Although aging affects the entire GM volume in the brain, as shown in **Supplementary Figure 5**, significant negative association between GM volume and age have been reported for several specific brain regions, i.e. a reduction in GM volume with age^9,10^. According to literature^9,10^ the insula, superior temporal areas and multiple gyri have shown significant age-related GM volume differences. However, due to the large size of most of these regions often only parts of these region were highlighted by the network. Interestingly, brain structures affected by age with higher p-value in literature^9,10^, were also more highlighted by the network, e.g.: caudate nucleus, amygdala, hippocampus and thalamus.

---

**Supplementary Table 1.** Characteristics of subjects with the 5-year age-stratified lowest quintile age gap values compared to the 5-year age-stratified highest quintile age gap values.

| Characteristic | Value lowest quintile (n=340) | Value highest quintile (n=350) | p-value |
| --- | --- | --- | --- |
| Age gap (years) | -5.7 ± 3.9 | 6.9 ± 4.5 | <0.001 |
| Grey matter volume (mL) | 605 ± 56.9 | 577.6 ± 56.2 | <0.001 |
| Systolic blood pressure (mmHg) | 138.9 ± 21.6 | 143.1 ± 21.0 | 0.009 |
| Mild cognitive impairment, n (%) | 15 (4.4) | 31 (8.9) | 0.013 |
| Diastolic blood pressure (mmHg) | 82.1 ± 10.8 | 84.1 ± 11.1 | 0.014 |
| Fasting glucose level (mmol/L) | 5.5 ± 1.2 | 5.7 ± 1.1 | 0.021 |
| Current or past smoker, n (%) | 102 (30.0) | 130 (37.1) | 0.027 |
| Body mass index (kg/m<sup>2</sup>) | 27.2 ± 3.9 | 27.8 ± 4.5 | 0.043 |
| Mini-Mental State Examination score | 28.0 ± 1.8 | 27.7 ± 2.1 | 0.095 |
| Total cholesterol (mmol/L) | 5.6 ± 1.0 | 5.5 ± 1.1 | 0.323 |
| APOEε4 carrier, n (%) | 92 (27.1) | 103 (29.4) | 0.418 |
| Female, n (%) | 187 (55.0) | 203 (58.0) | 0.428 |
| HDL cholesterol (mmol/L) | 1.4 ± 0.4 | 1.5 ± +- 0.4 | 0.549 |
| Age (years) | 65.5 ± 10.8 | 65.3 ± 11.0 | 0.771 |
| Years of education | 12.4 ± 3.8 | 12.3 ± 4 | 0.829 |
| Intracranial volume (mL) | 1465.8 ± 163.2 | 1466.3 ± 164.1 | 0.971 |

Values are presented in mean ± SD unless stated otherwise.
*Abbreviations:* number of participants (n); standard deviation (SD).

**Supplementary Figure 1.**
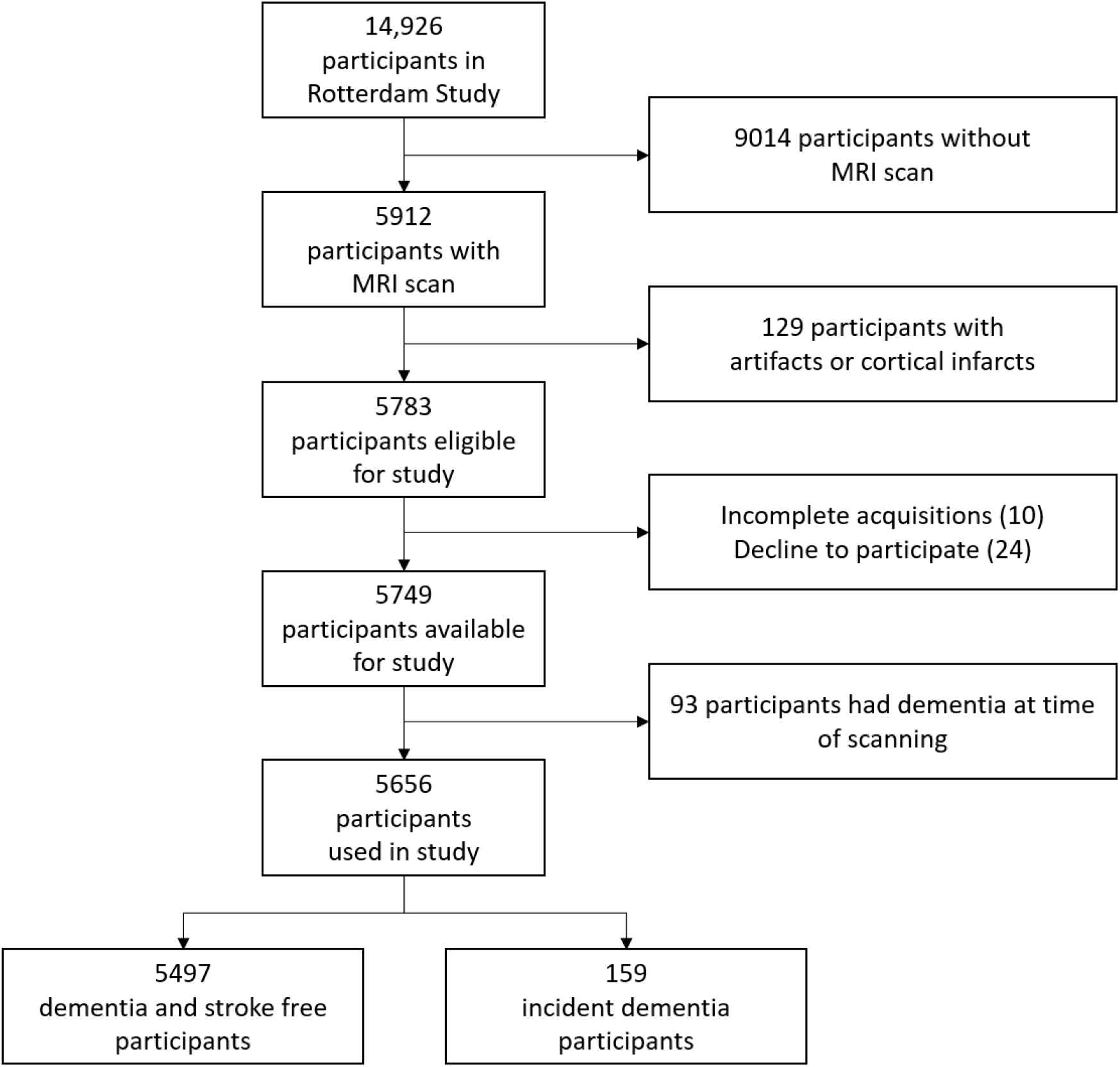
Flowchart showing the number of excluded participants per category.

**Supplementary Figure 2.**
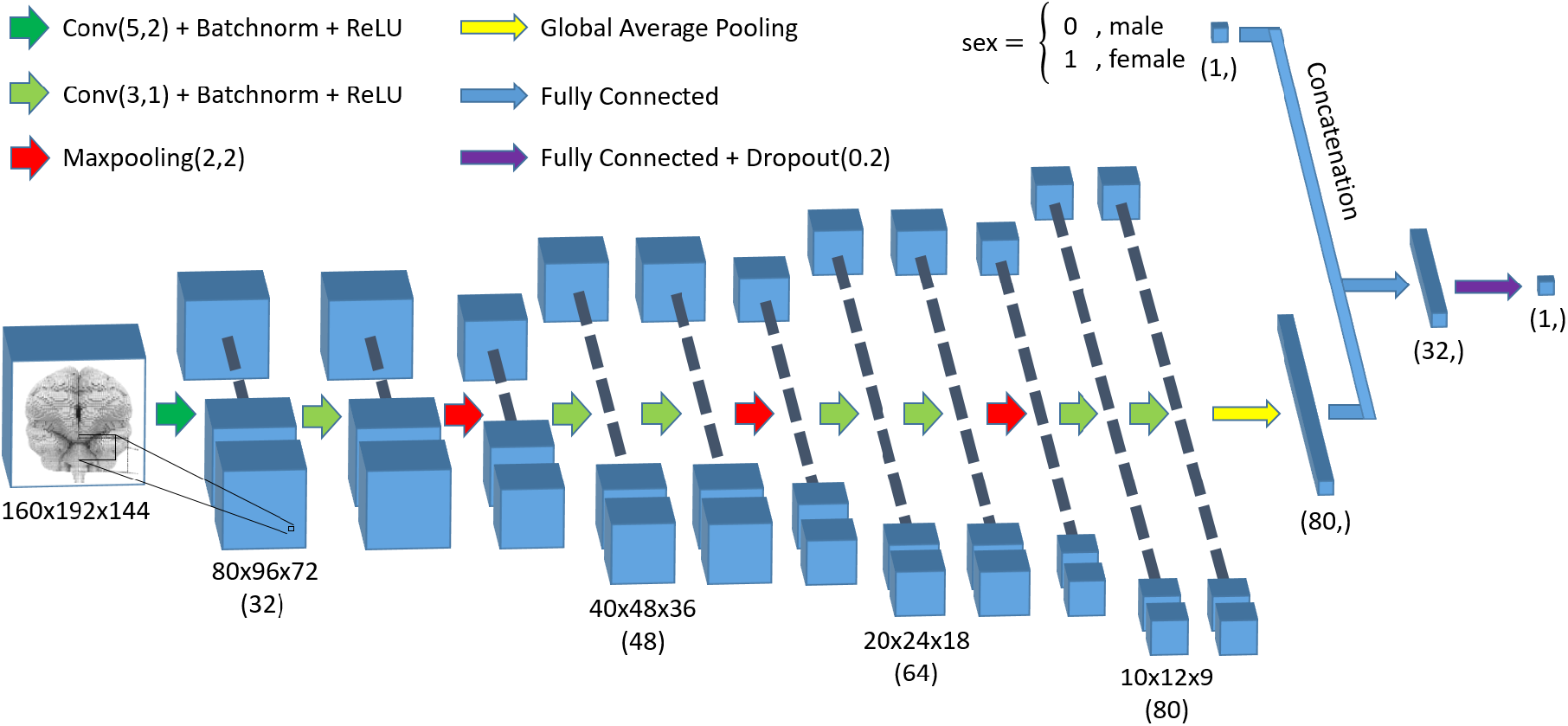
Graphical representation of the network architecture. The overall approach can be seen as four convolutional blocks ending on a pooling layer, which halves feature map dimensions. Hereafter, global average pooling extracts the final feature maps to a one-dimensional array of a single value per feature map. Fully connected layers are used to propagate to a single regression output. *Abbreviations:* kxkxk convolutional layer, with strides of sxsxs (CONV(k,s)); kxkxk max-pooling layer, with strides of sxsxs (Maxpooling(k,s)); batch normalization (Batchnorm); rectified linear unit (ReLU); dropout with probability p (Dropout(p)).

**Supplementary Figure 3.**
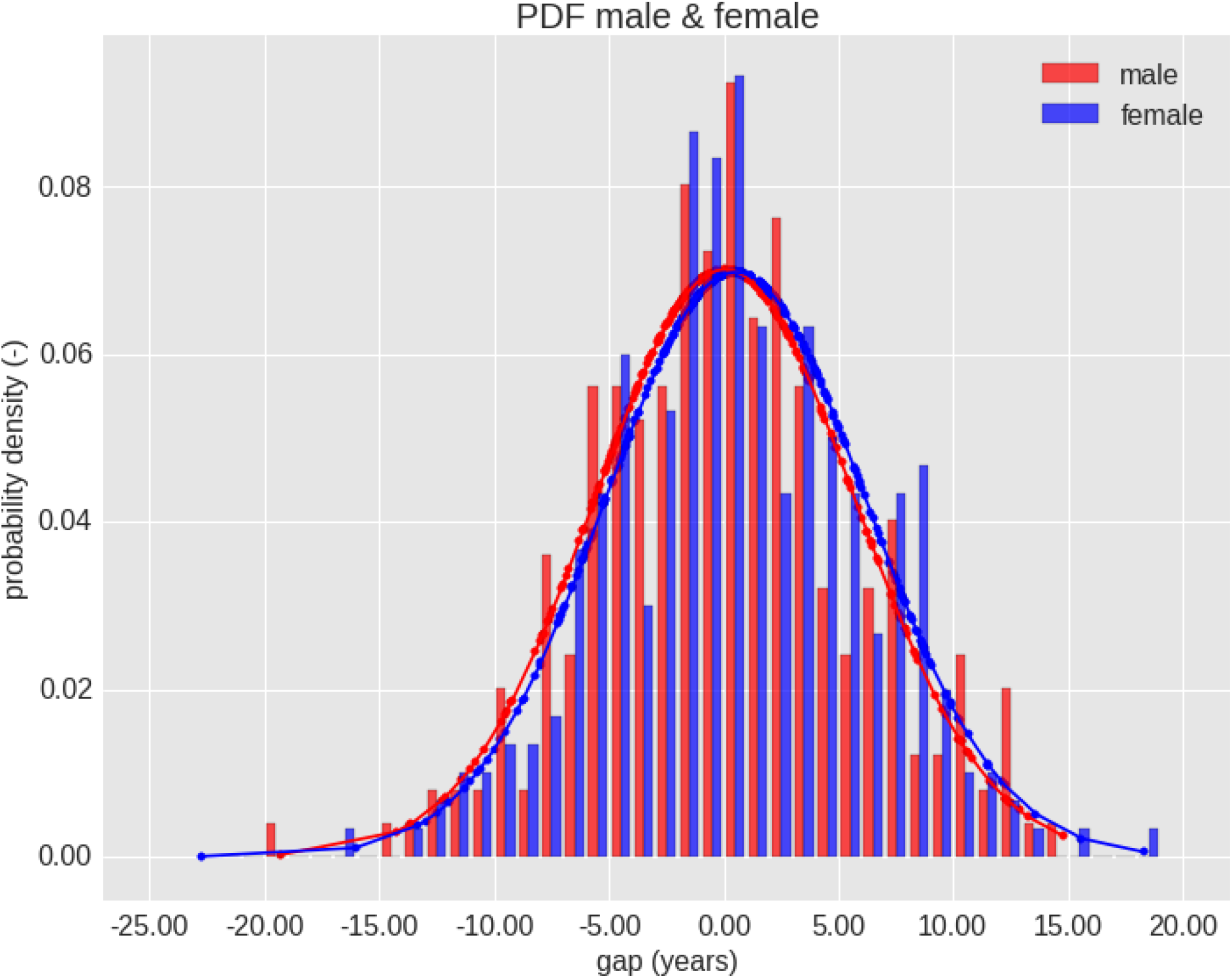
The probability density of the gap value (PAD) for male and female subjects. The distribution shows the difference in prediction for these two groups. Distributions are similar as mean η_female_ = 0.51 and variance σ^2^_female_ = 5.72 for female, whereas η_male_ = 0.04 and σ^2^_male_ = 5.69 for male. Resulting t-test showed no significant difference between the two groups as t(550) = −0.96 and p = 0.34.

**Supplementary Figure 4.**
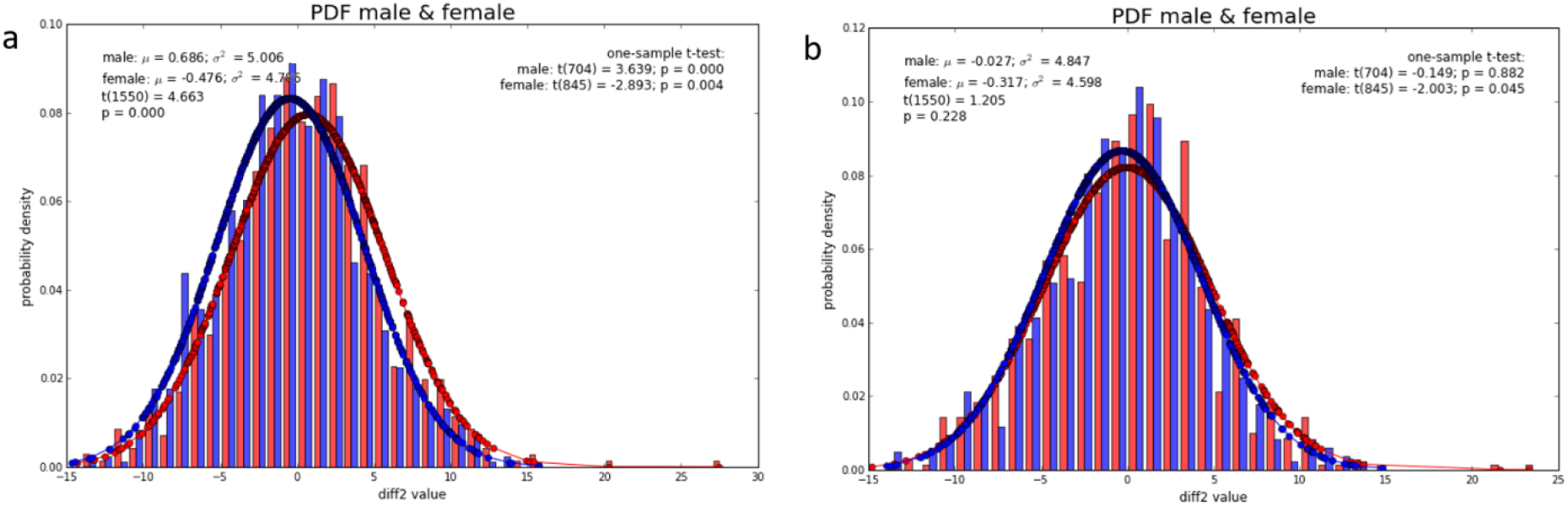
Effect of adding sex as a covariate to the model on the gap value distribution (red=male; blue=female). A comparison of the probability density functions for gap of two early trained models along with their respective t-test results. Both models have the exact same architecture with one the exception. a) Model uses only a single brain-MRI voxels input. b) Model uses two inputs, i.e. brain-MRI voxels and respective sex. Models were trained under the exact same settings.

**Supplementary Figure 5.**
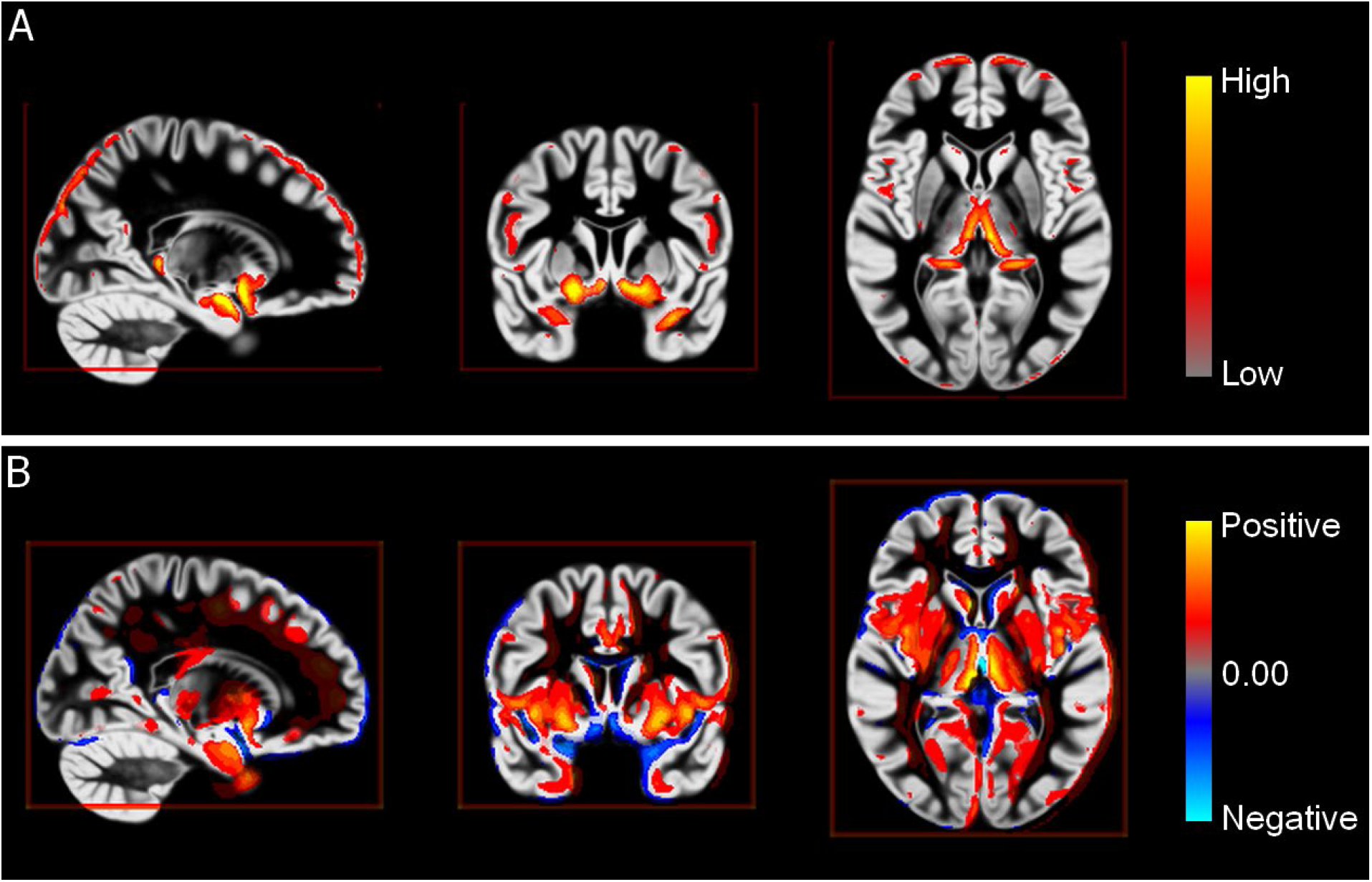
Grad-CAM attention map and attention map change overlaid on a brain template. **(A)** Grad-Cam attention map intensity per voxel. Voxel values in the attention map have been set at 0.65 minimum threshold and capped at 0.95 maximum to exclude background values and focus on more important regions. **(B)** Increase in attention map intensity over chronological age per voxel. Map include only voxels with a significant increase in voxel values (p<3e^−7^ after Bonferroni correction by number of GM voxels).

**Supplementary Figure 6.**
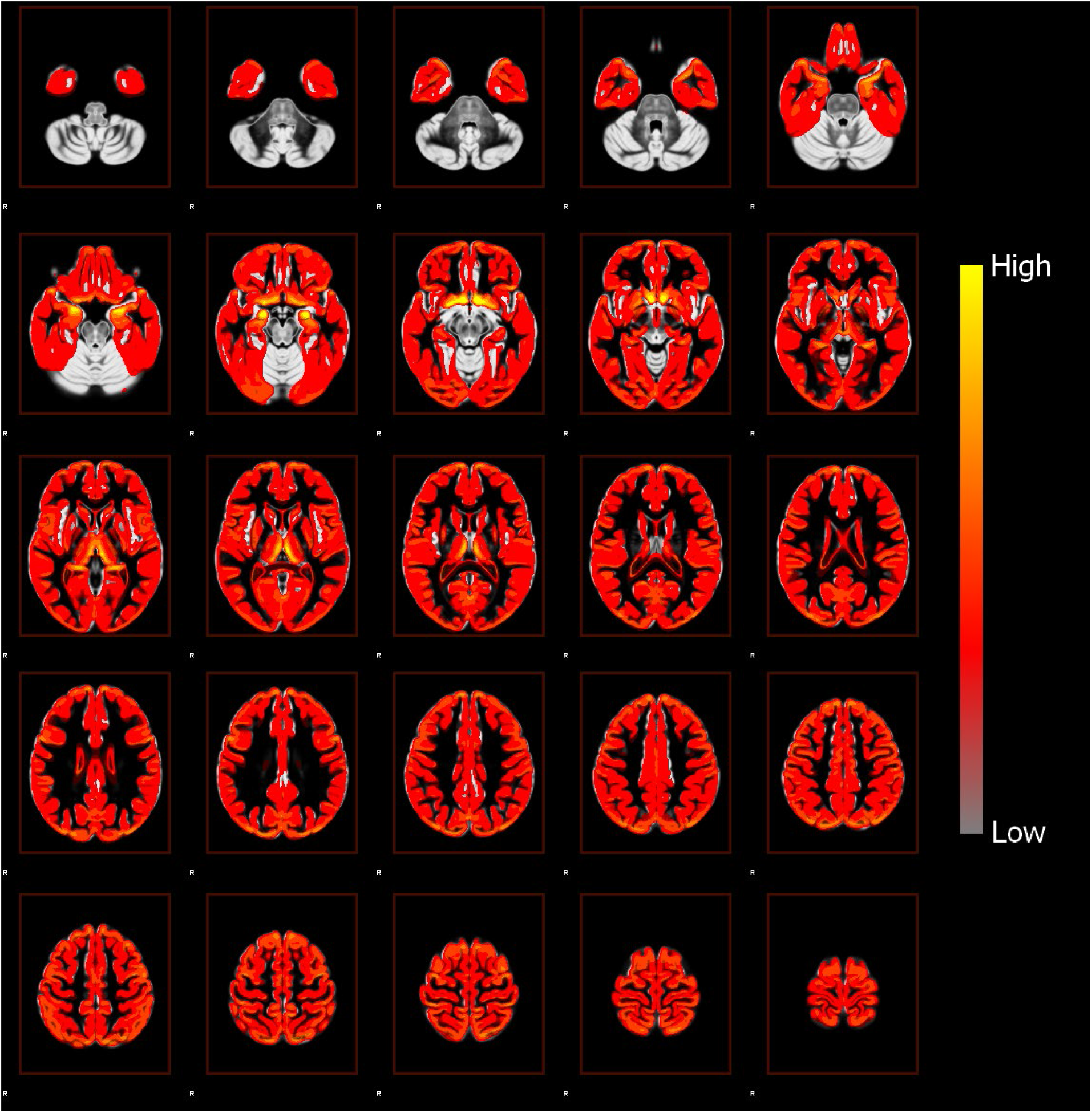
Grad-CAM attention map intensity per voxel overlaid on a brain template. Voxel values in the attention map have been set at 0.50 minimum and 1.00 maximum threshold to exclude background values and focus on more highlighted regions, according to normalization around 0.50 in the Grad-CAM implementation.

## REFERENCES

1. Vinke, E. J. et al. Trajectories of imaging markers in brain aging: The Rotterdam Study. Neurobiol. Aging (2018). doi:10.1016/j.neurobiolaging.2018.07.001

2. Manard, M., Bahri, M. A., Salmon, E. & Collette, F. Relationship between grey matter integrity and executive abilities in aging. Brain Res. 1642, 562–580 (2016).

3. Abbott, A. Dementia: A problem for our age. Nature 475, (2011).

4. Franke, K., Luders, E., May, A., Wilke, M. & Gaser, C. Brain maturation: Predicting individual BrainAGE in children and adolescents using structural MRI. Neuroimage 63, 1305–1312 (2012).

5. Liem, F. et al. Predicting brain-age from multimodal imaging data captures cognitive impairment. Neuroimage 148, 179–188 (2017).

6. Cole, J. H. et al. Predicting brain age with deep learning from raw imaging data results in a reliable and heritable biomarker. Neuroimage 163, 115–124 (2017).

7. Kaufmann, T. et al. Genetics of brain age suggest an overlap with common brain disorders. bioRxiv (2018). doi:10.1101/303164

8. Holmes, G. L., Milh, M. D. M. & Dulac, O. Maturation of the human brain and epilepsy. Handbook of Clinical Neurology 107, (2012).

9. Franke, K., Gaser, C., Manor, B. & Novak, V. Advanced BrainAGE in older adults with type 2 diabetes mellitus. Front. Aging Neurosci. 5, (2013).

10. Ikram, M. A. et al. The Rotterdam Scan Study: design update 2016 and main findings. Eur. J. Epidemiol. 30, 1299–1315 (2015).

11. Good, C. D. et al. A voxel-based morphometric study of ageing in 465 normal adult human brains. Neuroimage 14, 21–36 (2001).

12. Roshchupkin, G. V. et al. Fine-mapping the effects of Alzheimer’s disease risk loci on brain morphology. Neurobiol. Aging 48, 204–211 (2016).

13. Vrooman, H. A. et al. Multi-spectral brain tissue segmentation using automatically trained k-Nearest-Neighbor classification. Neuroimage 37, 71–81 (2007).

14. Smith, S. M. & Nichols, T. E. Threshold-free cluster enhancement: Addressing problems of smoothing, threshold dependence and localisation in cluster inference. Neuroimage 44, 83–98 (2009).

15. Mutlu, U. et al. Association of Retinal Neurodegeneration on Optical Coherence Tomography With Dementia. JAMA Neurol. 1–8 (2018). doi:10.1001/jamaneurol.2018.1563

16. McKhann, G. et al. Clinical diagnosis of Alzheimer’s disease. Neurology 34, 939 (1984).

17. Román, G. et al. Vascular dementia: diagnostic criteria for research studies. Neurology 43, 250–260 (1993).

18. LeCun, Y., Bottou, L., Bengio, Y. & Haffner, P. Gradient-based learning applied to document recognition. Proc. IEEE 86, 2278–2323 (1998).

19. Krizhevsky, A., Sutskever, I. & Hinton, G. E. ImageNet Classification with Deep Convolutional Neural Networks. Adv. Neural Inf. Process. Syst. 1–9 (2012). doi:http://dx.doi.org/10.1016/j.protcy.2014.09.007

20. Simonyan, K. & Zisserman, A. Very Deep Convolutional Networks for Large-Scale Image Recognition. arXiv Prepr. 1–10 (2014). doi:10.1016/j.infsof.2008.09.005

21. Selvaraju, R. R. et al. Grad-CAM: Visual Explanations from Deep Networks via Gradient-Based Localization. Proc. IEEE Int. Conf. Comput. Vis. 2017–Octob, 618–626 (2017).

22. Shrout, P. E. & Fleiss, J. L. Intraclass correlations: Uses in assessing rater reliability. Psychol. Bull. 86, 420–428 (1979).

23. Matsuda, H. Voxel-based Morphometry of Brain MRI in Normal Aging and Alzheimer’s Disease. Aging Dis. 4, 29–37 (2013).

24. Roses, A. D. & Saunders, A. M. APOE is a major susceptibility gene for Alzheimer’s disease. Curr. Opin. Biotechnol. 5, 663–667 (1994).

25. Rossum, G. Van & Drake, F. L. Python Reference Manual. Python Software Foundation (2001). Available at: http://www.python.org.

26. Ascher, D., Dubois, P., Hinsen, K., Hugunin, J. & Oliphant, T. Numerical Python. Lawrence Livermore National Laboratory (2001). Available at: http://www.pfdubois.com/numpy/.

27. Chollet, F. Keras. Github repository (2015). Available at: https://github.com/fchollet/keras.

28. R Core Team. R: A language and environment for statistical computing. R Foundation for Statistical Computing Available at: http://www.r-project.org/.

29. Aylward, E. H. et al. MRI volumes of the hippocampus and amygdala in adults with Down’s syndrome with and without dementia. Am. J. Psychiatry (1999). doi:10.1176/ajp.156.4.564

30. Wachinger, C., Salat, D. H., Weiner, M. & Reuter, M. Whole-brain analysis reveals increased neuroanatomical asymmetries in dementia for hippocampus and amygdala. Brain (2016). doi:10.1093/brain/aww243

## Supplementary references

1. Sathya R, Abraham A. Comparison of Supervised and Unsupervised Learning Algorithms for Pattern Classification. Int J Adv Res Artif Intell. 2013;2(2):34–38. doi:10.14569/IJARAI.2013.020206.

2. Xinghuo Yu, M. Onder Efe and OK. A General Backpropagation Algorithm for Feedforward Neural Networks Learning. IEEE Trans Neural Networks. 2002;13(1):251–254.

3. LeCun Y, Bottou L, Bengio Y, Haffner P. Gradient-based learning applied to document recognition. Proc IEEE. 1998;86(11):2278–2323. doi:10.1109/5.726791.

4. Krizhevsky A, Sutskever I, Hinton GE. ImageNet Classification with Deep Convolutional Neural Networks. Adv Neural Inf Process Syst. 2012:1–9. doi:http://dx.doi.org/10.1016/j.protcy.2014.09.007.

5. Xue-Wen Chen, Xiaotong Lin. Big Data Deep Learning: Challenges and Perspectives. IEEE Access. 2014;2:514–525. doi:10.1109/ACCESS.2014.2325029.

6. Işin A, Direkoglu C, Şah M. Review of MRI-based Brain Tumor Image Segmentation Using Deep Learning Methods. Procedia Comput Sci. 2016;102(August):317–324. doi:10.1016/j.procs.2016.09.407.

7. Ker J, Wang L, Rao J, Lim T. Deep Learning Applications in Medical Image Analysis. IEEE Access. 2018:1–1. doi:10.1109/ACCESS.2017.2788044.

8. Perez L, Wang J. The Effectiveness of Data Augmentation in Image Classification using Deep Learning. 2017. http://arxiv.org/abs/1712.04621.

9. Manard M, Bahri MA, Salmon E, Collette F. Relationship between grey matter integrity and executive abilities in aging. Brain Res. 2016;1642:562–580. doi:10.1016/j.brainres.2016.04.045.

10. Matsuda H. Voxel-based Morphometry of Brain MRI in Normal Aging and Alzheimer’s Disease. Aging Dis. 2013;4(1):29–37. http://www.pubmedcentral.nih.gov/articlerender.fcgi?artid=3570139&tool=pmcentrez&rendertype=abstract.

